# The Arabidopsis bZIP19 and bZIP23 transcription factors act as zinc-sensors to control plant zinc status

**DOI:** 10.1101/2020.06.29.177287

**Authors:** Grmay H. Lilay, Daniel P. Persson, Pedro Humberto Castro, Feixue Liao, Ross D. Alexander, Mark G.M. Aarts, Ana G.L. Assunção

## Abstract

Zinc (Zn) is an essential micronutrient for plants and animals because of its structural and catalytic roles in many proteins. Zn deficiency affects *ca*. two billion people, mainly those living on plant-based diets that rely on crops from Zn deficient soils. Plants maintain adequate Zn levels through tightly regulated Zn homeostasis mechanisms, involving Zn uptake, distribution and storage, but it was not known how they sense Zn status. We use *in vitro* and *in planta* approaches to show that the *Arabidopsis thaliana* F-group bZIP transcription factors bZIP19 and bZIP23, which are the central regulators of the Zn deficiency response, act as Zn sensors by binding Zn^2+^ ions to a Zn sensor motif (ZSM). Deletions or modifications of this ZSM disrupts Zn binding, leading to a constitutive transcriptional Zn deficiency response, which causes a significant increase in plant and seed Zn accumulation. Since the ZSM is highly conserved in F-bZIPs across land plants, the identification of the first plant Zn-sensor will promote new strategies to improve the Zn nutritional quality of plant-derived food and feed, and contribute to tackle the global Zn deficiency health problem.

## Main text

Zinc (Zn) is an essential micronutrient for plants and animals because of its structural and catalytic role in a large number of proteins as a cofactor in many enzymes, transcription factors and protein interaction domains (Andreini *et al*., 2006). Zinc deficient soils are widespread globally and Zn deficiency is estimated to affect about one-third of the world’s human population. It affects mainly populations living on plant-based diets that rely on crops from Zn deficient soils (Welch and Graham, 2004; Wessells and Brown, 2012).

To maintain adequate intracellular Zn levels, and avoid deficiency or toxic excess, plants rely on a tightly regulated Zn homeostasis network, which involves Zn uptake, transport, distribution and storage activities (Clemens, 2001; Colvin *et al*., 2010). One of the primary means by which cells regulate Zn homeostasis and Zn levels is through Zn-dependent changes in the expression of genes required for Zn uptake, transport and storage (Choi and Bird, 2014). The Zn sensing mechanisms and the corresponding Zn-dependent transcriptional regulation are being unravelled in prokaryotes, fungi and vertebrates (Patzer and Hantke, 1998; Andrews, 2001; Eide, 2009) but in plants, the mechanisms behind Zn-sensing remained elusive so far.

The *Arabidopsis thaliana* (Arabidopsis) bZIP19 and bZIP23 transcription factors are the central transcriptional regulators of the Zn deficiency response (Assunção *et al*., 2010). They belong to the F group of bZIP proteins (F-bZIP) and appear to be paralogs resulting from a duplication event associated with the Brassicaceae. Both contain a characteristic region with two short amino acid sequences rich in cysteine and histidine residues (Cys/His-rich motif), located at the N-terminus, which is highly conserved in F-bZIP homologs across land plants (Castro *et al*., 2017). bZIP19 and bZIP23 bind to *Zinc Deficiency Response Elements* (*ZDREs*) in the promoters of target genes that are activated under Zn deficiency (Assunção *et al*., 2010). These target genes comprise a small set of Zn homeostasis genes involved in Zn uptake, transport and distribution. Among them are genes encoding ZIP (Zinc-regulated/Iron-regulated Protein) family Zn transporters, which mediate Zn uptake into the cell, and NAS (Nicotianamine Synthase) enzymes that produce the Zn chelator nicotianamine (NA) involved in Zn distribution (Guerinot, 2000; Assunção *et al*., 2010; Clemens *et al*., 2013; Inaba *et al*., 2015). The *bzip19 bzip23* double mutant (*bzip19/23*) is hypersensitive to Zn deficiency, but with no visible phenotype under Zn sufficiency (Assunção *et al*., 2010). bZIP19 and bZIP23 are partially redundant, with the *bzip19* single mutant showing a mild Zn deficiency phenotype (Assunção *et al*., 2010; Inaba *et al*., 2015), possibly because the expression patterns of *bZIP19* and *bZIP23* do not exactly overlap (Lilay *et al*., 2019). The Zn-dependent activity of these transcription factors is controlled at the protein level, with repressed activity at Zn sufficiency (Lilay *et al*., 2019).

A theoretical model hypothesizes that the Zn-dependent activity of bZIP19 and bZIP23 can be directly modulated by Zn^2+^ ions, via direct binding of Zn to their Cys/His-rich motif (Assunção *et al*., 2013). Cys and His are Zn-coordinating amino acids characteristic of many Zn-binding proteins, including the classic Cys_2_His_2_ Zn-finger motif (Maret, 2013). To test whether the Cys/His-rich motif of bZIP19 and bZIP23 can bind Zn, we performed an *in vitro* Zn-protein binding assay with the bZIP19 and bZIP23 native proteins and a modified bZIP19 protein, of which the two Cys/His-rich sequences of the motif were deleted (bZIP19 del1 del2). The Cys/His-rich motif of the bZIP19 and bZIP23 proteins are nearly identical, with fully conserved Cys and His residues (Fig. 1A). The proteins were expressed and purified from *E. coli*, followed by incubation with the stable isotope ^67^Zn, and analysed by Size Exclusion Chromatography coupled to an Inductively Coupled Plasma Mass Spectrometer (SEC-ICP-MS). The sulphur (S) signal, serving as a proxy signal for Cys-and/or methionine-(Met) containing proteins (Persson *et al*., 2009), was analysed as the ^48^SO oxide product ion, and the Zn signal was monitored as the ^67^Zn and ^66^Zn isotopes. The *in vitro* Zn-protein binding assay revealed binding of Zn by the bZIP19 protein, as well as by the bZIP23 protein, based on the observed co-elution of the ^48^SO and ^67^Zn signals in the chromatogram after ∼90 s (Fig. 1B). The incubation with ^67^Zn and the ∼96% ^67^Zn/^66^Zn ratio in the peak at ∼90 s strongly indicates the binding of ^67^Zn to the protein. The bZIP19 del1 del2 protein also eluted after ∼90 s, as indicated by the ^48^SO signal, however it did not bind to Zn, neither to ^67^Zn nor to ^66^Zn (Fig. 1B). The S: ^67^Zn molar ratio for the bZIP19 and bZIP23 proteins was the same, suggesting that each protein binds two Zn atoms (Fig. 1C).

**Figure 1.**
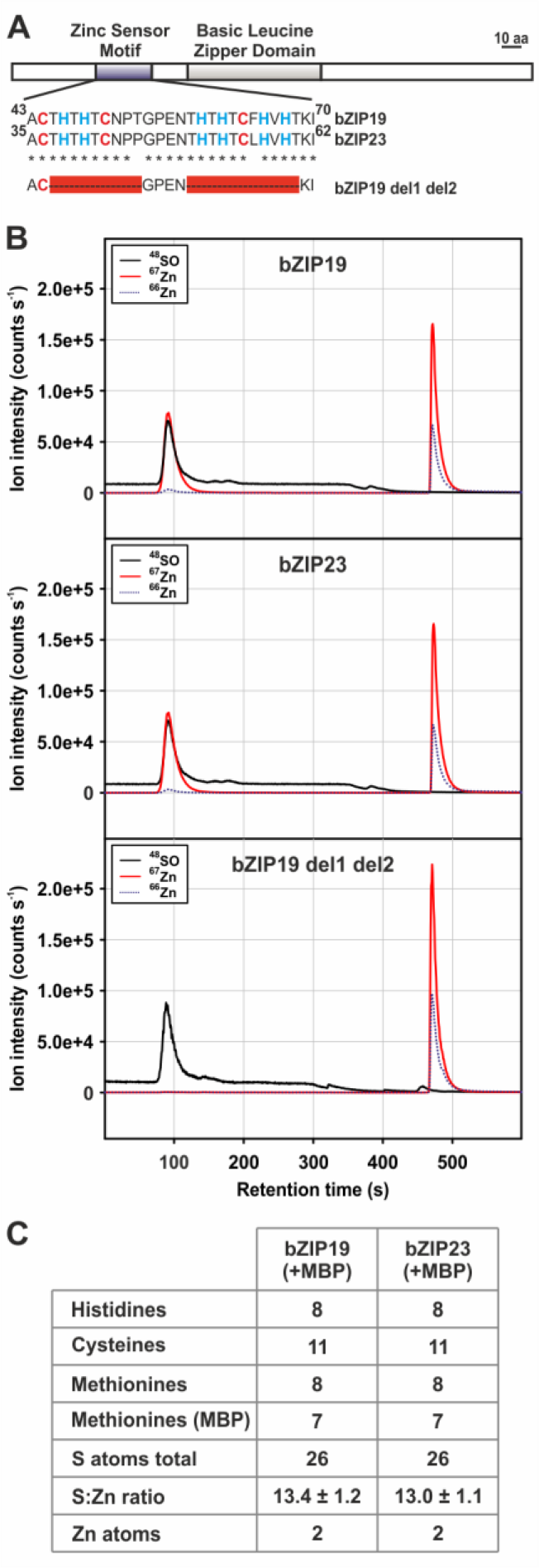
*In vitro* Zn-protein binding assay. (A) Alignment of the conserved Cys/His-rich Zn sensor motif in bZIP19 and bZIP23 proteins, and the deleted Cys/His-rich amino acid sequences of this motif in the bZIP19 del1 del2 mutant protein. Cys (C) and His (H) residues are represented in red and blue, respectively, and deletions are represented as red boxes. (B) SEC-ICP-MS chromatogram showing the speciation of Zn and S, run in O_2_ mode, in samples of purified proteins (*ca*. 15 μM) incubated with 30 μM of ^67^Zn stable isotope: Both the bZIP19 and bZIP23 native proteins (top and middle panels, respectively) show co-elution between the ^48^SO and ^67^Zn signals after ∼90 s, with an additional ^67^Zn peak eluting after ∼490 s representing surplus unbound Zn. The small ^66^Zn signals (eluting at ∼90 s and ∼490 s) indicate that the Zn binding to the protein originated from the ^67^Zn added at the incubation. bZIP19 del1 del2 protein (motif deletion mutant; bottom panel) shows no co-elution of the ^48^SO and ^67^Zn signals. Here, the late-eluting ^67^Zn peak (∼490 s) was more intense because no Zn had been bound to the protein. (C) The S: Zn molar ratio for the bZIP19 and bZIP23 was calculated from external calibration curves, generating a S: ^67^Zn ratio of 13.0-13.4 (*i*.*e*. 13 S atoms per Zn atom). The analysed bZIP19 and bZIP23 proteins have 26 S-containing amino acids (8 Met and 11 Cys, plus 7 Met in the maltose binding peptide (MBP) fusion protein). Thus, our measurements indicate that each protein binds to two Zn atoms. The control Zn-protein binding assay with MBP alone showed no Zn binding. Three independent protein extractions and 4-6 runs were analysed for each bZIP protein.

Having shown that bZIP19 and bZIP23 proteins bind Zn *in vitro* and that removal of the Cys/His-rich motif disrupts Zn binding, we next verified the significance of this result *in planta*. We used the Arabidopsis *bzip19/23* double mutant line, containing an Arabidopsis *ZIP4 promoter::GUS* fusion (*pZIP4::GUS*), as a platform for analysis. ZIP4 is a ZIP family Zn transporter and one of the target genes of bZIP19 and bZIP23, with *ZIP4* transcription induced at Zn deficiency (Assunção *et al*., 2010; Castro *et al*., 2017). The Zn deficiency hypersensitive phenotype of the *bzip19/23* mutant is complemented by the overexpression of bZIP19 or bZIP23, which restores the wild-type *ZIP4* expression pattern (Lilay *et al*., 2019). Hence, we used the line with *pZIP4::GUS* in the null *bzip19/23* background (*bzip19/23*-*pZIP4::GUS*), complemented by overexpression of bZIP19 motif mutant variants, to analyse the GUS expression as a proxy for the bZIP19 regulatory activity. The *bzip19/23*-*pZIP4::GUS* line was complemented either with the native bZIP19 or the bZIP19 del1 del2 (Fig. 2A), and the corresponding seedlings were exposed to Zn deficiency, sufficiency or excess, and subsequently analysed through histochemical GUS staining. As previously observed (Castro *et al*., 2017), the analysis of the *bzip19/23*-*pZIP4::GUS* line did not show GUS staining at any Zn supply condition, whereas the *pZIP4::GUS* line (Col-0 background) showed GUS staining at Zn deficiency that markedly decreased at Zn sufficiency, and was absent at Zn excess. The same pattern was observed in the *bzip19/23-pZIP4::GUS-bZIP19* line where a gradient of decreasing GUS staining was observed upon increasing Zn exposure (Fig. 2B). A similar decrease in target gene expression had also been observed for transcriptome data of Arabidopsis grown at deficiency, sufficiency or excess Zn for *ZIP4* and other *ZIP* transporter family target genes of bZIP19 and bZIP23 (Azevedo *et al*., 2016; Castro *et al*., 2017). In contrast, our analysis of the *bzip19/23-pZIP4::GUS-bZIP19 del1 del2* line showed a constitutive GUS staining, with no visible differences between Zn deficiency, sufficiency or excess (Fig. 2B), indicating that this mutated bZIP19 is still able to activate the *ZIP4* promoter, but is no longer responsive to changes in Zn status.

**Figure 2.**
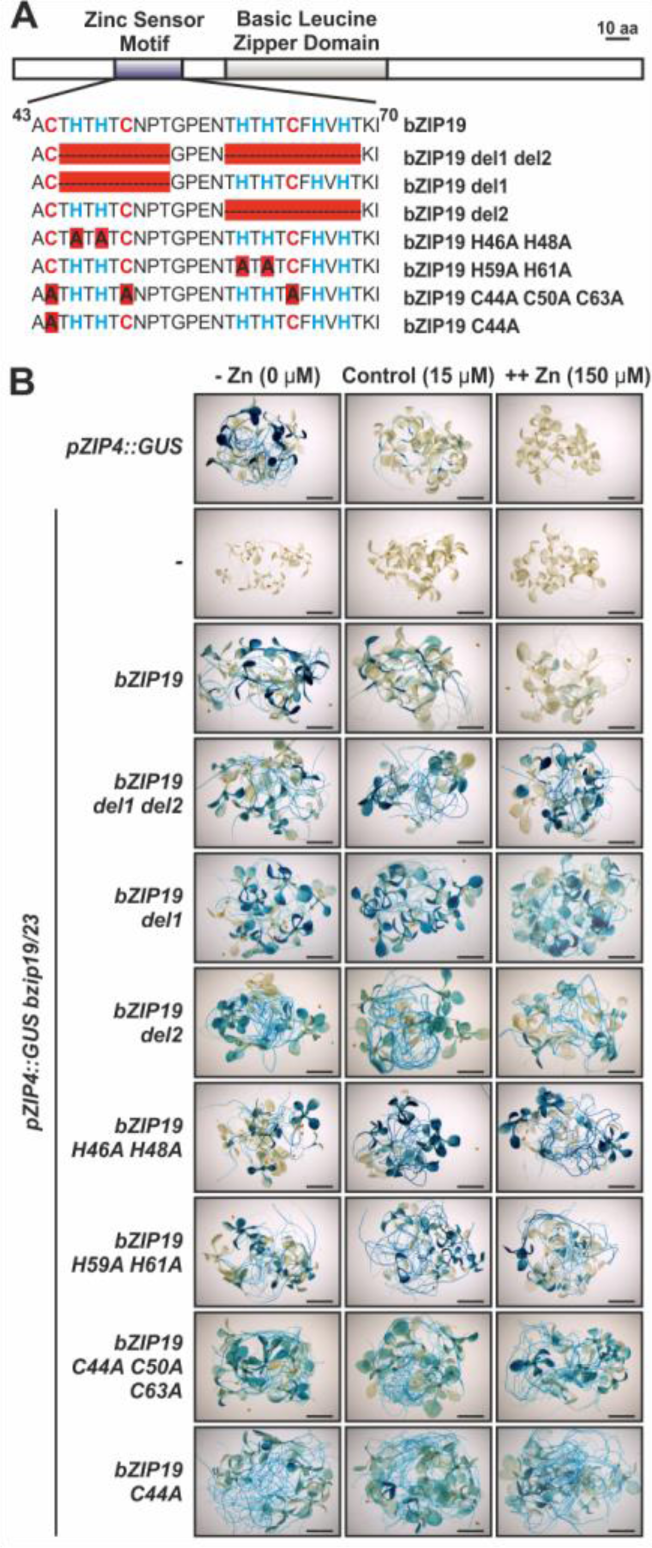
*In planta* analysis of the ability of bZIP19 Cys/His-rich Zn sensor motif variants to activate transcription of the *GUS* gene from the *ZIP4* promoter in a *bzip19/23*-*pZIP4::GUS* background. (A) Alignment of the Cys/His-rich motif in each bZIP19 variant. Cys (C) and His (H) residues are represented in red and blue, respectively. Deletion and single amino acid substitution variants are represented as red boxes. (B) Histochemical GUS staining analysis of *pZIP4::GUS, bzip19/23*-*pZIP4::GUS, bzip19/23*-*pZIP4::GUS-bZIP19* (native bZIP19), *bzip19/23*-*pZIP4::GUS-bZIP19 del1 del2*, and *bzip19/23*-*pZIP4::GUS-bZIP19 del1, -del2*, and, *-C44A C50A C63A, -C44A, -H46A H48A* and *-H59A H61A* variant lines. Images represent ten to fifteen 14-day-old seedlings grown on Zn deficient (-Zn), sufficient (control, 15 μM) or excess (++Zn, 150 μM) 1/2MS media. Six to ten independently transformed lines (T2) for each variant were analyzed.

We analysed additional bZIP19 motif variants, including deletions of only half the motif (bZIP19 del1 and bZIP19 del2) and amino acid substitutions of Cys (C) and His (H) with alanine (A), either substituting all C residues (bZIP19 C44A C50A C63A), one C residue only in the first half motif (bZIP19 C44A), or two H residues in each half motif (bZIP19 H46A H48A and bZIP19 H59A H61A) (Fig. 2A). Analysis of *bzip19/23*-*pZIP4::GUS* complemented with each of these bZIP19 motif variants showed the same GUS expression as the *bzip19/23-pZIP4::GUS-bZIP19 del1 del2* line, *i*.*e*. constitutive GUS staining with no visible differences between the different Zn supply concentrations (Fig .2B). These results show that both Cys/His-rich sequences in the motif, and even individual Cys or His residues, are needed for Zn deficiency-dependent activation of the *ZIP4* target gene by bZIP19. Considering that Cys and His are Zn-coordinating amino acids (Maret, 2013), our results indicate that all Cys and His residues within the Cys/His-rich motif are necessary to provide the adequate structural environment for Zn-protein binding. Although further investigation is needed at the level of protein structure and its coordination with Zn^2+^ ions (*i*.*e*. the Zn binding sites within the Cys/His-rich motif and binding affinity), it is likely that the Zn-protein binding causes a conformational change that affects the regulatory activity of bZIP19 and bZIP23. Such a conformational change could interference with the DNA-binding of the transcription factors, with their dimerization process or other protein-protein interactions, or with protein stability (Assunção *et al*., 2013).

In addition to the analysis of the *ZIP4* promoter GUS expression (Fig. 2B), we also quantified the actual expression level of three target genes (*ZIP4, ZIP5* and *NAS2*) of bZIP19 and bZIP23. These analyses showed the characteristic Zn deficiency-induced target gene expression in the wild-type and in the complemented *bzip19/23-bZIP19* line (Lilay *et al*., 2019), and constitutive target gene activation in the *bzip19/23-bZIP19 del1 del2* variant line (Fig. 3A-C), in agreement with the GUS expression analysis (Fig. 2B). Next, we investigated how this Zn-independent target gene activation impacts plant Zn content. Element analysis of 6-week-old plants grown hydroponically with control Zn supply revealed that, while the shoot Zn concentrations of the wild-type, the *bzip19/23* mutant and the complemented *bzip19/23-bZIP19* plants are not significantly different, the Zn concentrations in shoots of the bZIP19 del1 del2 plants are significantly higher (around 2-to 3-fold) relative to the wild-type (Fig. 3D). The profiling of other elements (iron (Fe), copper (Cu), manganese (Mn) and phosphorous (P)) showed some significant differences between lines, but mostly irrespective of the genotype, and much less pronounced than for Zn (Fig. S1). This indicates that the deregulation of the Zn deficiency response in plants has a largely Zn-specific Zn accumulation effect, which offers promising perspectives for Zn biofortification.

**Figure 3.**
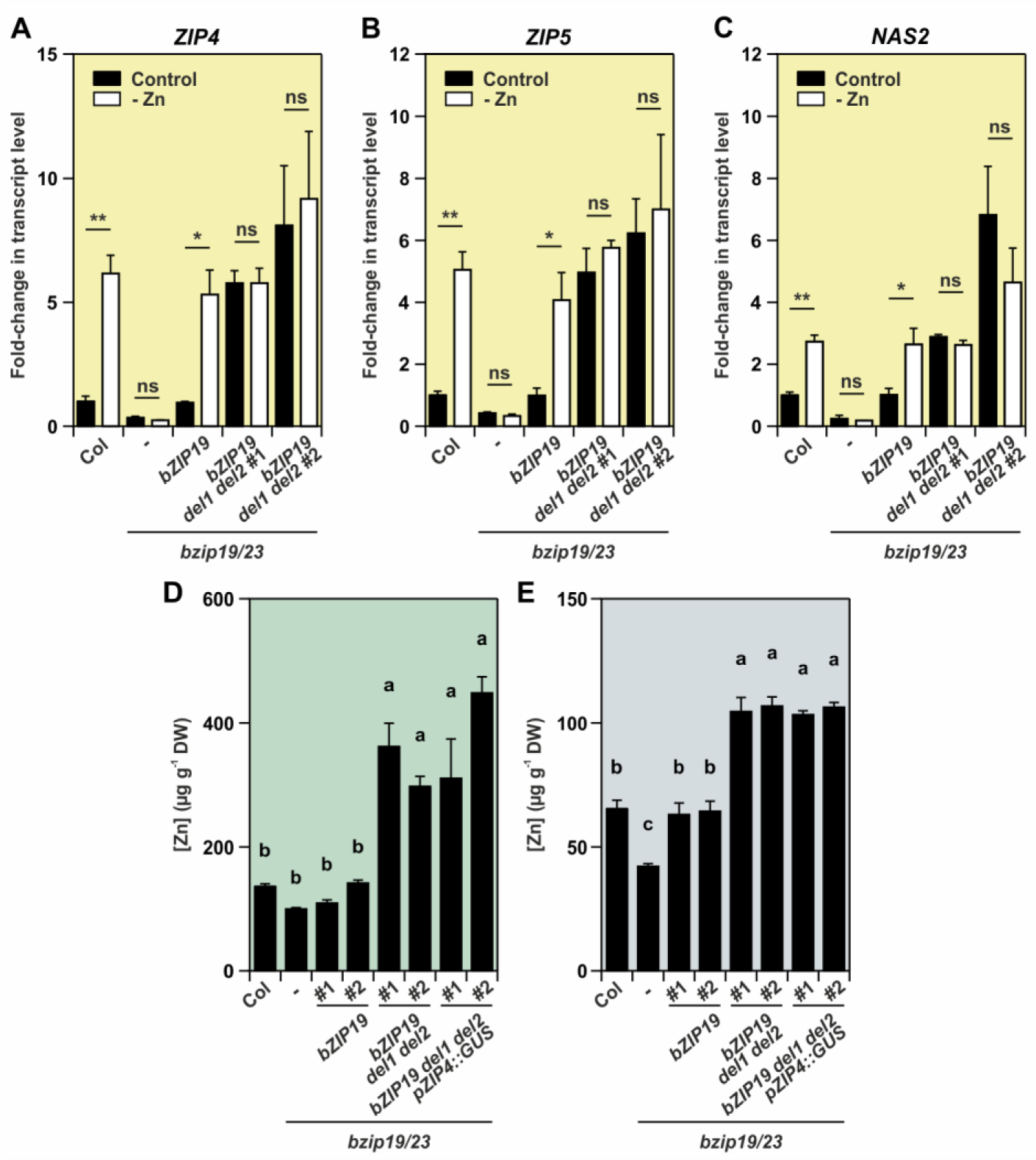
Effects on gene expression (A-C) and Zn concentration (D-E) upon expression of the bZIP19 del1 del2 variant in the *bzip19/23* and *bzip19/23*-*pZIP4::GUS* backgrounds. (A-C) Gene expression analysis of bZIP19 and bZIP23 target genes: *ZIP4, ZIP5* and *NAS2*, in 14-day-old seedlings of wild-type (Col), *bzip19/23* (-), *bzip19/23-bZIP19* and *bzip19/23-bZIP19 del1 del2* lines grown with control (dark bars) or Zn-deficient (–Zn, open bars) 1/2MS media. Bars represent mean fold-change in transcript level of three biological replica ±SE. Statistically significant differences between control and –Zn were determined by Student t-test (* p < 0.05, ** p < 0.01, *** p < 0.001, ns means not significant). (D-E) Average Zn concentration in shoots (D, n=4) of 6-week-old plants of wild-type (Col), *bzip19/23* (-), *bzip19/23-bZIP19, bzip19/23-bZIP19 del1 del2* and *bzip19/23*-*pZIP4::GUS-bZIP19 del1 del2* lines grown in hydroponics with control nutrient solution, and seeds (E, n=6) from the same lines harvested from soil-grown single plants. Different letters indicate significant differences (p < 0.05) after one-way ANOVA followed by Tukey’s post-hoc test. # indicates that homozygous T3 progeny of two independently transformed lines are used for each transgene.

We also investigated the ionome profile of the seeds and found that seeds of the bZIP19 del1 del2 lines had a significantly higher Zn concentration than wild-type seeds, with *ca*. 50% increase relative to the wild type (Fig. 3E). Again, the extent of the increase in Zn concentration, when compared to other elements (Fig. S2), indicates a largely Zn-specific effect.

We showed that Arabidopsis bZIP19 and bZIP23 proteins bind Zn^2+^ ions *in vitro*, which is disrupted by removal of the conserved Cys/His-rich motif. We also showed that the Zn deficiency transcriptional activation of their target genes, *in planta*, depends on the Cys/His-rich motif being intact, including all the individual Cys and His residues. In addition, this Zn deficiency transcriptional activation is Zn specific, since it is not affected by deficiency or sufficiency of other metal micronutrients but is only repressed by Zn sufficiency (Fig. S3). Together, these results provide robust evidence that bZIP19 and bZIP23 act as Zn sensors, through direct binding of Zn^2+^ to their Cys/His-rich motif, and we propose referring to this Cys/His-rich motif as Zn-sensor motif (ZSM).

The Zn sensing mechanism conferred by the ZSM appears to provide plants with a one-step link between the cellular Zn status and the transcriptional Zn-deficiency response (Fig. 4). The Zn deficiency response of eukaryotic cells is best studied in *Saccharomyces cerevisiae* yeast, where the Zap1 Zn-responsive transcriptional activator is the central regulator of Zn homeostasis (Zhao *et al*., 1998). In this case, the Zap1-mediated regulatory activity involves sensing cellular Zn status through complex and independent mechanisms of Zn binding to two different activation domains and the DNA binding domain of Zap1 (Lyons *et al*., 2000; Bird *et al*., 2003; Frey *et al*., 2011). In plants, although knowledge on other micronutrient homeostasis regulation has progressed significantly, especially for Fe (Castro *et al*., 2018; Kobayashi, 2019; Kim *et al*., 2019), their sensor proteins and molecular mechanisms through which plants perceive micronutrient deficiency remains unidentified. For example, Fe deficiency-inducible genes involved in Fe uptake and distribution are regulated by sophisticated transcriptional networks that comprise several transcription factors. Namely, Fe uptake is mediated by IRT1 transporter, a ZIP family member, that in addition to being transcriptionally induced by Fe-deficiency, is also known to have a non-Fe metal sensing mechanism that drives its own degradation. This is a post-translational regulatory mechanism of IRT1 that senses and protects against soil metal excess (Dubeaux *et al*., 2018). Since many of the transcription factors involved in Fe homeostasis regulation are transcriptionally induced in response to Fe deficiency, there should be other transcription enhancing factor, as well as Fe sensing and signalling molecules, acting upstream of these regulators (Kobayashi, 2019). Therefore, the Arabidopsis bZIP19 and bZIP23 proteins that we studied here are the first micronutrient sensor proteins upstream of a transcriptional regulatory network identified in plants.

**Figure 4.**
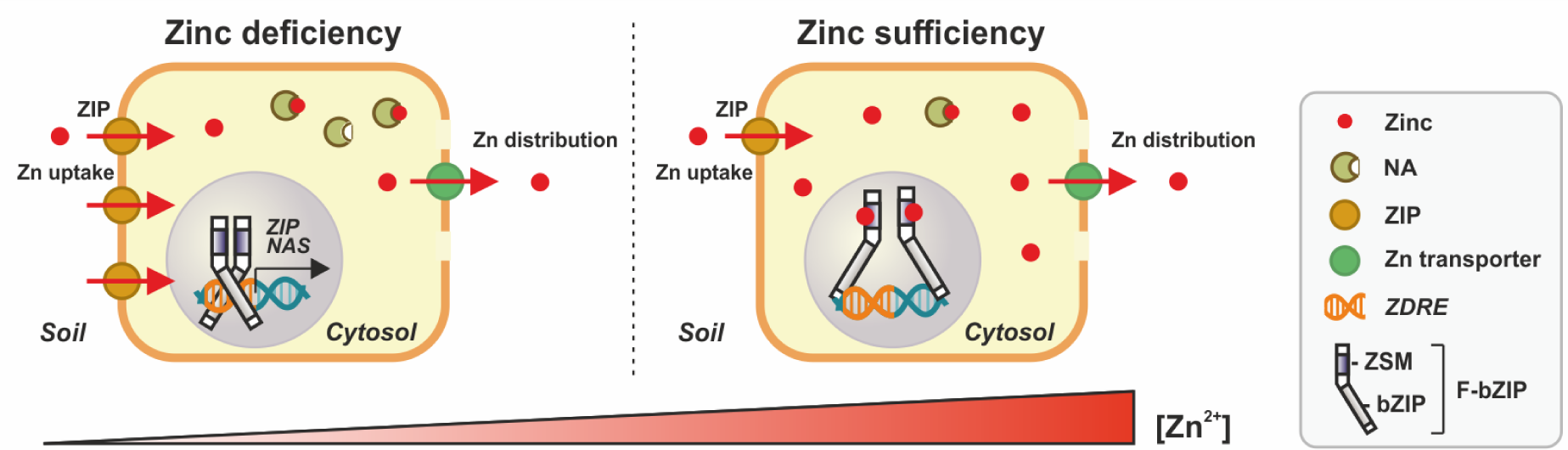
Schematic model of cellular Zn status sensing in Arabidopsis. bZIP19 and bZIP23 transcription factors, the central regulators of the Zn deficiency response, are depicted as F-bZIP. Under Zn deficiency, bZIP19 and bZIP23 dimerizing transcription factors, bind to the *Zn Deficiency Response Elements* (*ZDREs*) in the promoter of their target genes, activating their transcription. These include *ZIP* and *NAS* genes that contribute to Zn uptake and distribution. Under Zn sufficiency, cytosolic free or loosely bound Zn ions bind to the characteristic Cys/His-rich motif Zn sensor motif (ZSM) of bZIP19 and bZIP23 and this Zn-protein binding represses activity of the transcription factors (Assunção *et al*., 2013).

Human Zn malnutrition mainly affects populations relying on cereal grains as staple food, thus biofortification of seeds is a key challenge (White and Broadley, 2009; Cakmak and Kutman, 2018). Our finding of a 50% increase in seed Zn concentrations at sufficient Zn supply, without any apparent adverse effects on plant growth and development (Fig. S4), indicates a promising strategy for developing crops with enhanced Zn accumulation in seeds. The F-bZIP homologs are highly conserved across land plants, and the suggested conserved function as regulators of the Zn deficiency response (Castro *et al*., 2017), is supported by functional analysis of F-bZIP members from cereal crops (Evens *et al*., 2017; Nazri *et al*., 2017; Lilay *et al*., 2020). This work shows that individual Cys or His residues in the ZSM can be targeted to induce Zn-independent target gene expression. The bZIP19 and bZIP23 are the central regulators of the Arabidopsis Zn deficiency response that likely evolved to face fluctuations in soil Zn availability (Castro *et al*., 2017), and further research is needed on how disruption of their Zn-sensor ability to modulate Zn acquisition affects plant Zn homeostasis. Nonetheless, based on our results it is plausible that the activity of F-bZIP homologs can be modulated to improve Zn acquisition in seed crops by, for example, selection of naturally-occurring ZSM variants, or by CRISPR-Cas9-mediated precise editing technology (Shan *et al*., 2013),. This will be a welcome addition to the existing breeding efforts to address the problems of Zn deficiency in crops and plant-based human diets.

## Materials and Methods

### Plant material and growth conditions

The Arabidopsis genotypes used in this study were the wild-type accession Columbia (Col-0), the double T-DNA insertion mutant *bzip19 bzip23* (*bzip19/23*) obtained from a cross between homozygous *bzip19-1* (SALK_144252) and *bzip23-1* (SALK_045200) lines in Col-0 background, as described by (Assunção *et al*., 2010), and the *bzip19/23*-*pZIP4::GUS* line selected after a cross between the *bzip19/23* double mutant and a stable Col-0 transformant harbouring a construct with the Arabidopsis *ZIP4* promoter fused to the reporter gene *GUS* (*pZIP4::GUS)*, as described by (Castro *et al*., 2017). For agar-grown seedlings, sterilized seeds were sown on half-strength (1/2) MS medium containing 15 µM ZnSO_4_ (Zn sufficiency, control), no added Zn (Zn deficiency, -Zn), or 150 µM ZnSO_4_ (Zn excess, ++Zn). For the other micronutrient deficiencies, Fe (-Fe), Cu (-Cu) or Mn (-Mn) was omitted from the media. For hydroponically-grown plants, seeds were germinated and grown for 6 weeks with a modified half-strength Hoagland nutrient solution with either 2 µM ZnSO_4_ (Zn sufficiency, control) or with 0.002 µM ZnSO_4_ (Zn deficiency, -Zn), as described by (Lilay *et al*., 2019). Plants grown on the hydroponics setup or on the MS plates were in a growth chamber with 8/16 h light/dark cycle, with 125 µmol m^−2^ s^−1^ white light, 22/20 °C light/dark temperature, and 70% relative humidity. The analysed seeds were harvested from single plants of T3 homozygous lines grown on 0.4 L pots with a mixture of peat (80%) and vermiculite (20%) in a growth chamber with the above mentioned conditions, except a light/dark cycle of 16/8 h. Four to six pools of seeds were obtained for each genotype.

### Plasmid construction and plant transformation

To generate the *bzip19/23-bZIP19* and *bzip19/23*-*pZIP4::GUS-bZIP19* lines, the full-length protein coding sequence of *bZIP19* (At4g35040) was amplified from Arabidopsis (Col-0) cDNA. The cloning into pEarleyGate-102 Gateway vector (Earley *et al*., 2006) carrying a *CaMV35S* promoter, a C-terminal CFP and HA-tag to generate the *pCaMV35S::bZIP19-CFP-HA* construct, is described by (Lilay *et al*., 2019). To generate the bZIP19 Cys/His-rich motif mutant variants, primers were designed to introduce different deletions and amino acid substitutions by PCR in the native bZIP19 (Table S1). The mutant variants were cloned to generate *pCaMV35S::bZIP19mut-CFP-HA* constructs, as described above. The transformation into *Agrobacterium tumefaciens* cells and Arabidopsis were performed as described by (Lilay *et al*., 2019). The constructs corresponding to bZIP19, bZIP19 del1 del2, bZIP19 del1, bZIP19 del2, bZIP19 C44A C50A C63A, bZIP19 C44A, bZIP19 H46A H48A and bZIP19 H59A H61A were transformed into the Arabidopsis *bzip19/23*-*pZIP4::GUS* line, and the constructs corresponding to bZIP19 and bZIP19 del1 del2 were transformed into the Arabidopsis *bzip19/23* double mutant. The *bzip19/23-bZIP19* line is the same as the previously reported *bzip19/23-OE19* line (Lilay *et al*., 2019). Transgenic plants were screened for Basta (phosphinothricin) resistance and T2 generation seeds of six to ten independently transformed lines per construct were selected, and further homozygous T3 generation seeds of three to four lines were selected. To generate the protein expression constructs, the *pCaMV35S::bZIP19-CFP-HA, pCaMV35S::bZIP19 del1 del2-CFP-HA* and *pCaMV35S::bZIP23-CFP-HA* (Lilay *et al*., 2019) constructs were amplified using forward and reverse primers containing *Xmn*I and *Hind*III restriction sites, respectively (Table S1), followed by PCR product purification (PureLink PCR Purification Kit, Invitrogen) and cloning into the *Xmn*I*/Hind*III restriction sites of pMAL-c2 vector, which contains an IPTG-inducible *tac* promoter and an N-terminal Maltose Binding Protein (MBP) tag. The resulting constructs were verified by sequencing and were transformed into *E. coli* Rosetta 2 strain competent cells for heterologous expression.

### Histochemical staining for B-glucuronidase (GUS) assay and imaging

Histochemical GUS staining was performed with 14-day-old seedlings of *bpZIP4::GUS, bzip19/23*-*pZIP4::GUS, bzip19/23*-*pZIP4::GUS-bZIP19* and *bzip19/23*-*pZIP4::GUS-bZIP19* motif variant lines, grown on control, –Zn or ++Zn 1/2MS media. Six to ten independently transformed T2 generation seedlings were tested for each line. Seedlings were immersed in GUS staining solution containing 50 mM phosphate buffer, 10 mM Na_2_-EDTA, 20% (v/v) methanol, 0.1% (v/v) Triton X-100, 1.4 mM K3[Fe(CN)6], 1.4 mM K4[Fe(CN)6].3H2O with 1.9 mM X-Gluc and were incubated overnight at 37°C in the dark (Jefferson *et al*., 1987).

After incubation, the pigments were removed by repeated incubations in 50%, 70% and 96% (v/v) ethanol, and seedlings were stored in 70% (v/v) glycerol. Bright field images of GUS-stained seedlings, *ca*. ten to fifteen per independent line, were recorded with a Leica M205FA Stereo Fluorescence Microscope. All experiments were consistent between independent lines and within bZIP19 motif variant line. One representative image was used to construct Fig 2B. For the analysis of the specificity of the Zn deficiency response, 12-day-old seedlings of *pZIP4::GUS* and *bzip19/23*-*pZIP4::GUS* lines grown on control, -Zn, -Fe, -Cu or –Mn 1/2MS media were analysed as described above.

### Protein extraction and *in vitro* Zn-binding assay

The *E. coli* Rosetta 2 clones with *pMAL-bZIP19, pMAL-bZIP19 del1 del2* mutant, and *pMAL-bZIP23* were cultured on LB medium supplemented with selective antibiotics for 2–3 hours at 37°C, until OD_600_ of 0.6, then 0.1 M IPTG was added and bacterial cultures were grown for 2 more hours and centrifuged at 4,000 g for 20 min at 4°C. The pellet was suspended in extraction buffer (20 mM Tris-HCl pH 7.4, 200 mM NaCl, 1 mM EDTA, 1 mM DTT) supplemented with Complete Protease Inhibitor Cocktail (Roche). Cells were disrupted by sonication, centrifuged at 14,000 *g* for 30 min and soluble protein fraction was purified using MBPTrap Columns, according to manufacturer’s instructions (GE Healthcare Bio-Sciences). Protein concentration was measured with a Nanodrop spectrophotometer (Thermo-Scientific), and protein integrity and expected molecular weight were verified with a protein gel electrophoresis (SDS-PAGE) stained with Coomassie Blue (Fig. S5). For the *in vitro* Zn-binding assay, 15 μM of the fusion proteins were incubated with 10 mM DTT for 30 min at room temperature, followed by buffer exchange with 50 mM Tris-HCl, pH 7.2, and incubation with 30 μM of ^67^Zn stable isotope (67-Zn metal, 94.8% enriched, Trace Sciences International Corp., Richmond Hill, ON, Canada) for 90 min at room temperature. The samples were then analyzed by Size Exclusion Chromatography coupled to an Inductively Coupled Plasma Mass Spectrometer (SEC-ICP-MS), monitoring the ^48^SO, ^66^Zn and ^67^Zn signals. The SEC column was a Biobasic SEC 300 (300 × 7.8; 5 µm particle size, 300 Å pore size; Thermo Fisher Scientific, Waltham, MA, USA), mounted with a pre-column in front (Biobasic SEC 300 pre-column; Thermo Fisher Scientific). The HPLC (Dionex model UltiMate 3000; (Thermo Fisher Scientific), equipped with a DAD detector. All connections were of PEEK material with an internal diameter of 150 μm (nanoViper, Dionex; Thermo Fisher Scientific). The outlet from the DAD detector was connected to an ICP-MS (Agilent 8800 ICP-QQQ-MS; Agilent Technologies) operated in oxygen mode. Hence, S was analyzed as the oxide product ions (^48^SO^+^), using the S signal as a proxy signal for protein. Zn was analyzed as their parent ions. Each experiment was repeated with three independent protein extractions.

### Real-time quantitative RT-PCR analysis

Fourteen-day-old seedlings of Arabidopsis wild-type (Col-0), *bzip19/23, bzip19/23-bZIP19* and *bzip19/23-bZIP19 del1 del2* lines, grown on control or -Zn 1/2MS media, were harvested and immediately frozen in liquid nitrogen in pools of 5 seedlings per line and per Zn treatment x 3 different plates grown simultaneously and considered as biological replica. Two independently transformed T3 homozygous lines of *bzip19/23-bZIP19 del1 del2* (#1, #2) were analyzed. Total RNA extraction and cDNA synthesis were performed as described by (Lilay *et al*., 2019). Primers were designed using NCBI Primer-BLAST (www.ncbi.nlm.nih.gov/tools/primer-blast) and the primer amplification efficiency for each primer pair was between 1.9 and 2.1 (Table S1). RT-qPCR was performed with a LightCycler 96 Real-Time PCR System (Roche Diagnostics), using HOT FIREPol EvaGreen qPCR Mix (Solis BioDyne) in a 20 µL PCR reaction mixture, as described by (Lilay *et al*., 2019). The Arabidopsis *Actin-2* (*ACT2*, At3g18780) was used as reference gene. Reactions were performed in three technical replicas per biological replica and in 3 biological replicas per line and Zn treatment. The calculated cycle threshold (Ct) value for each gene was normalized to the reference gene calculated Ct value. The relative transcript levels were expressed against the wild-type grown with control conditions, and calculated according to the Livak 2^-ΔΔCT^ method (Livak and Schmittgen, 2001).

### Tissue element analysis

The rosette from 6-week-old hydroponically grown plants of Arabidopsis wild-type (Col-0), *bzip19/23, bzip19/23-bZIP19, bzip19/23-bZIP19 del1 del2* and *bzip19/23*-*pZIP4::GUS-bZIP19 del1 del2* lines, grown with control nutrient solution, were harvested (4-6 rosette/plants per line). Two independently transformed T3 homozygous lines of *bzip19/23*-*bZIP19 del1 del2* (#1, #2) and *bzip19/23*-*pZIP4::GUS-bZIP19 del1 del2* (#1, #2) were analyzed. Tissue digestion was performed with ultra-pure acids (70% HNO_3_ and 15% H_2_O_2_) at 240°C and 80 bars for 15 min in a pressurized microwave oven (Ultrawave, Milestone Inc.) enabling the amount of acid and final dilution to be adjusted according to the amount of plant material (Olsen *et al*., 2016). All samples were diluted to 3.5 % HNO_3_ prior to multi-elemental analysis using Inductively-Coupled Plasma Optical Emission Spectrometry (ICP-OES) (5100 ICP-OES, Agilent Technologies). The ICP-OES was equipped with a SeaSpray nebuliser and a double pass scott type spray chamber. Moreover, automatic sample introduction was performed from a ASX-520, CETAC auto-sampler from 15 mL falcon tubes. Certified reference material (NIST1515, apple leaf, National Institute of Standards and Technology, USA) was included to evaluate data quality and drift correction was performed based on drift samples for every 20 samples. Data were processed using Agilent ICP Expert Software version 7.3. Seeds of each line (*ca*. 15 mg x 6 replica) were also analysed following the same procedure.

### Statistical analysis

To compare lines or treatments we used one-way ANOVA followed by Tukey’s post-hoc test, or Student’s t-test, as appropriate, calculated with IBM SPSS Statistics V22.0 software.

## Acknowledgements

This work was supported by the Independent Research Fund Denmark, DFF-YDUN-program (4093–00245B); Portuguese Foundation for Science and Technology, FCT-IF program (IF/01641/2014); Novo Nordisk Foundation, Biotechnology-based Synthesis and Production Research program (NNF18OC0034598) [G.H.L., P.H.C., F.L., A.G.L.A.]; DFF-FTP-program (50544600-1126521001-112652) [D.P.P.] and Netherlands Genome Initiative (40-41009-98-11084) [R.A., M.M.G.A.].

## Author contributions

G.H.L., D.P.P., P.H.C., F.L., R.A. and A.G.L.A conducted experiments. G.H.L., D.P.P., P.H.C. and A.G.L.A conceived the study. G.H.L., M.G.M.A. and A.G.L.A wrote the manuscript. All authors reviewed and approved the manuscript.

## Competing interests

The authors declare no competing interests.

## Supplementary Material available

**Figure S1**. Shoot ionome analysis of the motif deletion mutant lines (bZIP19 del1 del2).

**Figure S2**. Seed ionome analysis of the motif deletion mutant lines (bZIP19 del1 del2).

**Figure S3**. Histochemical GUS staining analysis under different micronutrient deficiencies.

**Figure S4**. Deletion mutant lines grown with Zn deficient (-Zn) or sufficient (control) media.

**Figure S5**. Protein gel electrophoresis of purified bZIP19 and bZIP19 del1 del2 proteins.

**Table S1**. List of primers used in this study.

## Notes

### Competing Interest Statement

The authors have declared no competing interest.

