## Supplementary Material for "The Arabidopsis bZIP19 and bZIP23 transcription factors act as zinc-sensors to control plant zinc status"

**Supplementary Metarial includes:**

**Figure S1.** Shoot ionome analysis of the motif deletion mutant lines (bZIP19 del1 del2).
**Figure S2.** Seed ionome analysis of the motif deletion mutant lines (bZIP19 del1 del2).
**Figure S3.** Histochemical GUS staining analysis under different micronutrient deficiencies.
**Figure S4.** Deletion mutant lines grown with Zn deficient (-Zn) or sufficient (control) media.
**Figure S5.** Protein gel electrophoresis of purified bZIP19 and bZIP19 del1 del2 proteins.
**Table S1.** List of primers used in this study.

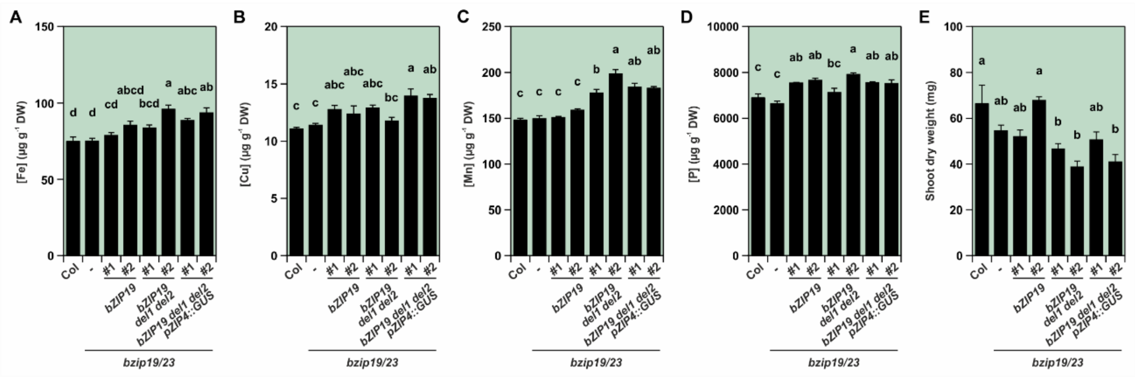


**Figure S1.** Shoot ionome analysis of the motif deletion mutant lines (bZIP19 del1 del2). Six-week-old plants of wild-type (Col), *bzip19/23* (-), *bzip19/23-bZIP19*, *bzip19/23-bZIP19 del1 del2* and *bzip19/23-pZIP4::GUS-bZIP19 del1 del2* lines grown in hydroponics with control nutrient solution. (A) Fe (B) Cu (C) Mn and (D) P concentration, and (E) dry weight (DW), of shoot tissue. # represents independently transformed T3 homozygous lines. Bars represent element concentration or DW of shoots as mean ±SE (n>4 plants). Different letters indicate significant differences ($p < 0.05$) after one-way ANOVA followed by Tukey's post-hoc test.

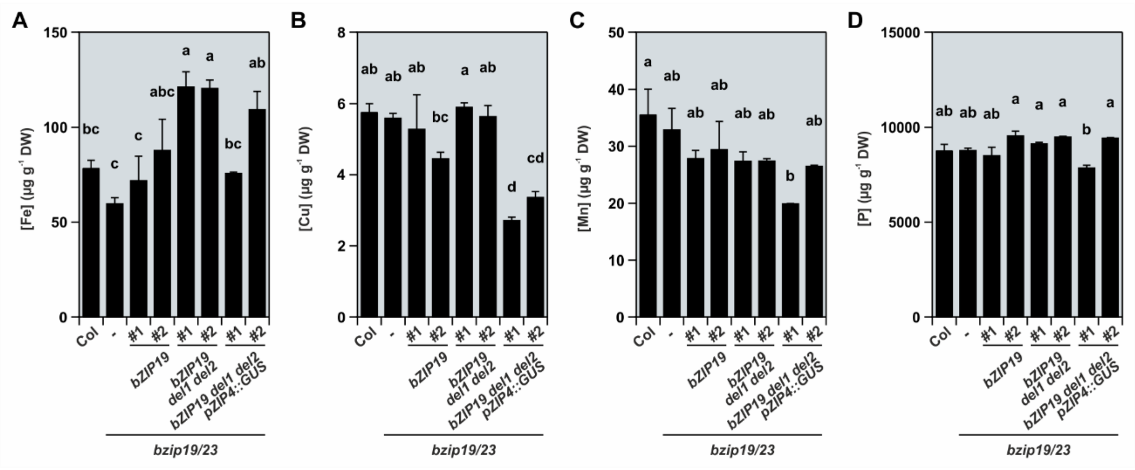


**Figure S2.** Seed ionome analysis of the motif deletion mutant lines (bZIP19 del1 del2). Seeds from soil-grown plants of wild-type (Col), *bzip19/23* (-), *bzip19/23-bZIP19*, *bzip19/23-bZIP19 del1 del2* and *bzip19/23-pZIP4::GUS-bZIP19 del1 del2* lines were analyzed. (A) Fe (B) Cu (C) Mn and (D) P concentration in seeds. # represents independently transformed T3 homozygous lines. Bars represent element concentration in seeds as mean ±SE (n=6). Different letters indicate significant differences ($p < 0.05$) after one-way ANOVA followed by Tukey's post-hoc test.

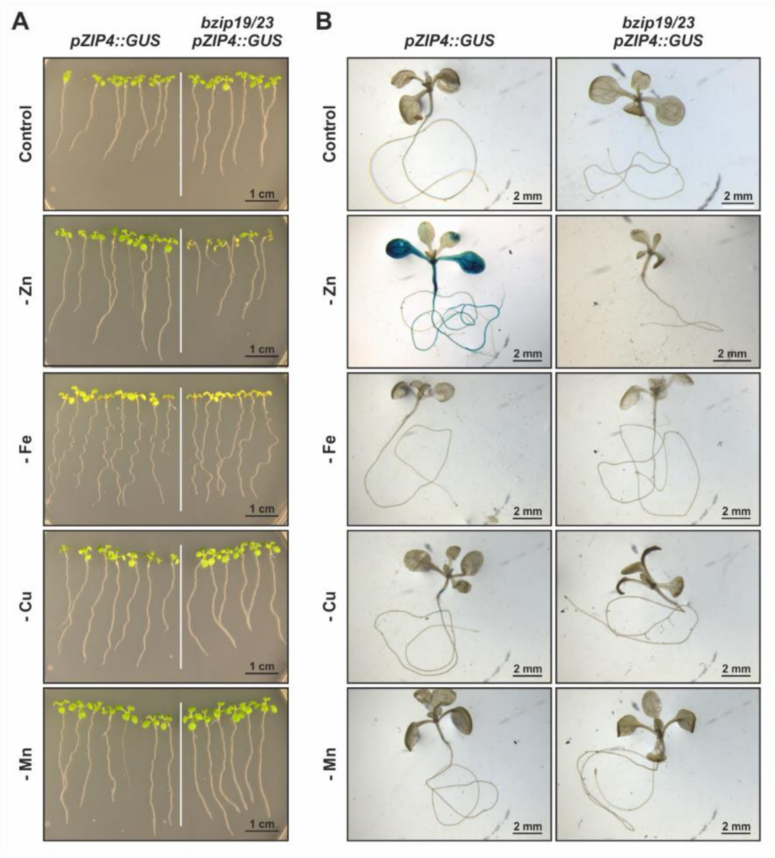


**Figure S3.** (A) Phenotypic analysis and (B) histochemical GUS staining of 12-day-old seedlings of *pZIP4::GUS* and *bzip19/23-pZIP4::GUS* lines grown on ½MS media with control or under different micronutrient deficiencies, *i.e.* Zn (-Zn), iron (-Fe), cupper (-Cu) and manganese (-Mn). Four to six plates (A), and 3-5 seedlings (B) per treatment and genotype were analyzed.

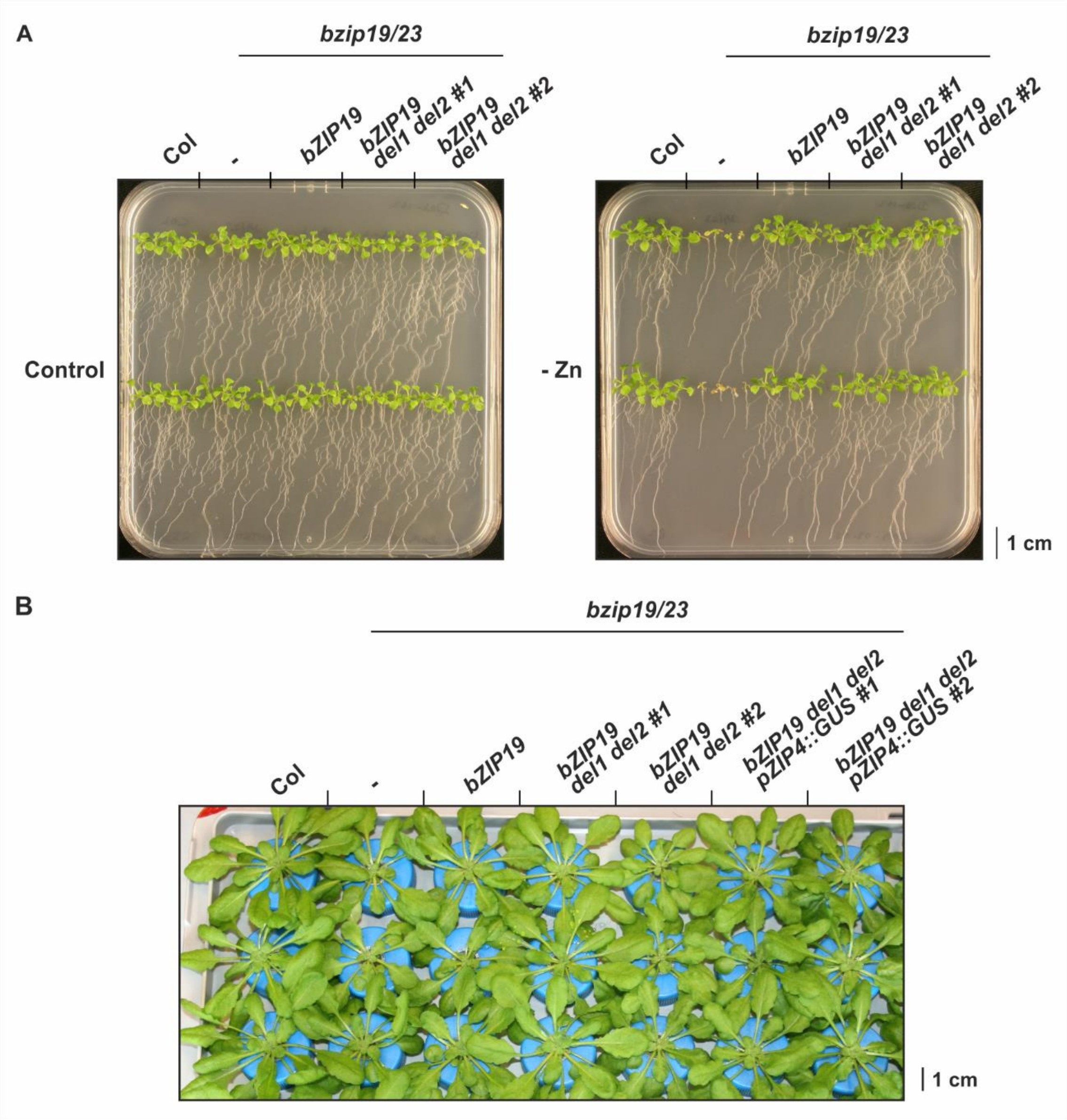


**Figure S4.** (A) Seedlings of 14-day-old Arabidopsis wild-type (Col), *bzip19/23* double mutant (-), *bzip19/23-bZIP19* and *bzip19/23-bZIP19 del1 del2* lines, grown with control or Zn deficiency (-Zn) ½MS media. (B) Six-week-old plants of wild-type (Col), *bzip19/23* (-), *bzip19/23-bZIP19*, *bzip19/23-bZIP19 del1 del2* and *bzip19/23-pZIP4::GUS-bZIP19 del1 del2* lines grown in hydroponics with control nutrient solution. # represents independently transformed T3 homozygous lines.

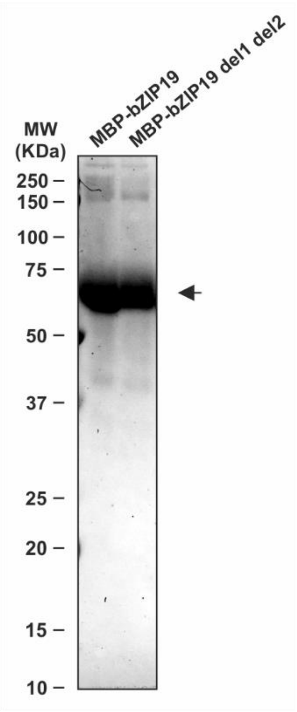


**Figure S5.** Protein gel electrophoresis (SDS-PAGE) stained with Coomassie Blue with 40 µg of the purified proteins from the *E. coli* Rosetta 2 clones with *pMAL-bZIP19* and *pMAL-bZIP19 del1 del2* mutant (*pMAL-bZIP23* not shown). The purified proteins show the expected molecular weight (arrow) *ca.* 68 KDa (28 KDa from bZIP19 + 40 KDa from MBP). Molecular weight markers (MW) are displayed.

**Table S1**. List of forward (F) and reverse (R) primers used in this study.

**Site-directed Mutagenesis**

| **Primer Name** | **Primer Sequence (5'→3')** |
|---|---|
| bZIP19 del1 F | TGCTGCTTGTGGACCAGAGAACACTCATAC |
| bZIP19 del1 R | TCTCTGGTCCACAAGCAGCATTAGTATCCA |
| bZIP19 del2 F | ACCAGAGAACAAGATTCTCCCGGATGAGAG |
| bZIP19 del2 R | GGAGAATCTTGTTCTCTGGTCCAGTGGGGT |
| bZIP19 H46A H48A F | GCTTGTACCGCCACTGCCACCTGTAACCCC |
| bZIP19 H46A H48A R | GGGGTTACAGGTGGCAGTGGCGGTACAAGC |
| bZIP19 H59A H61A F | GGACCAGAGAACACTGCTACTGCCACGTGC |
| bZIP19 H59A H61A R | GCACGTGGCAGTAGCAGTGTTCTCTGGTCC |
| bZIP19 C44A F | GATACTAATGCTGCTGCTACCCACACTCACACC |
| bZIP19 C44A R | GGTGTGAGTGTGGGTAGCAGCAGCATTAGTATC |
| bZIP19 C50A F | GCTTGTACCCACACTCACACCGCTAACCCC |
| bZIP19 C50A R | GGGGTTAGCGGTGTGAGTGTGGGTACAAGC |
| bZIP19 C63A F | CACTCATACTCACACGGCCTTCCATGTCCACAC |
| bZIP19 C63A R | GTGTGGACATGGAAGGCCGTGTGAGTATGAGTG |

Cloning into pMAL-c2

| **Primer Name** | **Primer Sequence (5'→3')** |
|---|---|
| bZIP19 F | CTATAGGAAGGATTTCAGTAATGGAAGACGGTGAGCTTGAT |
| bZIP19 R | CTTCAAAGCTTGATGATGCTTCAAACTGCTCTTGATGCACG |
| bZIP23 F | GATTAGGAAGGATTTCAGTAATGGACGACGGTGAGCTTGAG |
| bZIP23 R | CATCAAAGCTTACTTCAAACTGCTTTCGCTGCTCGAGGCTC |

RT-qPCR

| **Primer Name** | **Primer Sequence (5'→3')** |
|---|---|
| ZIP4 F | GATCTTCGTCGATGTTCTTTGG |
| ZIP4 R | TGAGAGGTATGGCTACACCAGCAGC |
| ZIP5 F | CGGGATTGTTGGCGTGGAAT |
| ZIP5 R | CCAAGACCCTCGAAGCATTG |
| NAS2 F | CGACGTGGTTAATTCGGTGG |
| NAS2 R | CGCGTGGACCTTAGAGCAAT |
| ACT2 F | CTAAGCTCTCAAGATCAAAGGCTTA |
| ACT2 R | ACTAAAACGCAAAACGAAAGCGGTT |
